# Protracted prediction: Neurodevelopment of reward processing in the adolescent cerebellum

**DOI:** 10.64898/2026.05.27.726906

**Authors:** Teagan S. Mullins, Jeremy Hogeveen

**Affiliations:** Institute for Graduate Clinical Psychology, Widener University.; Psychology Clinical Neuroscience Center, University of New Mexico.; Department of Psychology, University of New Mexico.

**Keywords:** Adolescent Neurodevelopment, Cerebellum, Reward Processing, Reinforcement Learning, Prediction Error, fMRI Meta-Analysis

## Abstract

Adolescence is characterized by heightened reward sensitivity and risky decision-making. Prevailing neurodevelopmental frameworks attribute these behavioral trends to a maturational imbalance between rapidly developing motivational brain regions, and slower-maturing prefrontal cognitive control circuitry. However, these models overlook the cerebellum—a structure that demonstrates protracted development across adolescence and reciprocal connectivity with both striatum and prefrontal cortex. Here, we conducted a systematic literature search and activation likelihood estimation (ALE) meta-analysis of functional magnetic resonance imaging (fMRI) studies examining reward anticipation and feedback processing in healthy adolescents (19 studies; 55 cerebellar peaks). In adults, a recent ALE analysis found a diffuse set of cognitive and affective cerebellar regions is recruited during reward anticipation, while feedback-locked activation is circumscribed and mostly restricted to the Vermis (Kruithof et al., 2023). In adolescents, we found evidence for a reward phase-specific spatial dissociation relative to adults. In adolescents, reward anticipation-locked activity is restricted to regions of the Vermis that are functionally connected to occipital and salience networks. Conversely, feedback processing elicits diffuse activation across bilateral Crus I/II regions that are connected to frontoparietal, default-mode, and limbic networks. These results suggest a *proactive* engagement of cortico-cerebellar networks during reward anticipation in adults, and a *reactive* recruitment of cortico-cerebellar networks post-feedback in adolescents. Adolescents may have less mature internal models for optimizing decision-making, and as a result they may require more widespread recruitment of cortico-cerebellar cognitive and affective circuitry post-feedback. Prevailing maturational imbalance models of youth decision-making should be expanded to incorporate developmental changes in cerebellar reward processing.

## Introduction

Adolescence is associated with a heightened sensitivity to rewards. Adolescent fMRI studies suggest this increased sensitivity to rewards is driven by enhanced recruitment of motivational brain regions including ventral striatum, amygdala, and orbitofrontal cortex (Cohen et al., 2010; Ernst et al., 2005; Galván and McGlennen, 2013; May et al., 2004). In parallel, cognitive control circuits anchored in lateral prefrontal cortex (PFC) undergo protracted development in adolescence, and are delayed in establishing the long-range connections with motivational regions critical for regulating motivated behavior (Crone and Steinbeis, 2017; Luna, 2009; Luna et al., 2010). Accordingly, ‘*dual systems models*’ of neurodevelopment have argued that an imbalance between hyperactivation of motivational brain regions in response to rewards, alongside hypoactivation of cognitive control circuits, drives the canonical increase in emotional and risky decision-making in adolescence (Casey et al., 2008; Fareri et al., 2008; Luna et al., 2015; Somerville et al., 2010; Steinberg, 2007). While these models have offered a critical heuristic for understanding adolescence, they fail to consider the potential relevance of another region that undergoes protracted maturation and plays a functional role in reward processing— the cerebellum.

Despite being one of the first structures to emerge in fetal development, the cerebellum is among the last brain regions to fully mature, a protracted developmental trajectory necessitated by the establishment of its complex reciprocal cerebellar-cerebral circuit anatomy (Dobbing, 1974). Similar to the prefrontal cortex, cerebellar structural volume follows an inverted ‘U-shaped’ developmental trajectory (Tiemeier et al., 2010). This volumetric growth occurs later than in much of the neocortex, with total cerebellar volume peaking between 12 and 16 years of age (≈12 years in females, ≈16 years in males; (Tiemeier et al., 2010)). Importantly, this maturation is not spatially uniform, and ‘cognitive’ regions of the cerebellum including Crus I-II appear to develop more slowly than more anterior sensorimotor areas (Bernard et al., 2016, 2015). Subsequently, the cerebellum’s pruning and shrinkage developmental trajectory is accelerated relative to the prefrontal cortex—while the PFC continues to undergo significant synaptic pruning throughout early adulthood (Petanjek et al., 2011), cerebellar grey matter volumes remain largely stabilized after the late teens / early 20s (Bernard and Seidler, 2014). As a result, it is hypothesized that the cerebellum may help to scaffold neurodevelopmental processes in connected cortical structures, by shaping activity-dependent myelination and cortical pruning in remotely connected regions like the PFC (Moberget et al., 2015; Wang et al., 2014).

In adulthood, the cerebellum likely plays a more central role in cognitive control and reward-guided behavior than traditionally realized. Recent evidence suggests the cerebellum functions as a critical modulatory hub that bridges motivational and control circuitry. It forms reciprocal loops with the prefrontal cortex which support high-level cognitive control (Buckner et al., 2011; Schmahmann, 2019), while simultaneously maintaining reciprocal connectivity with the basal ganglia (Bostan et al., 2010). Furthermore, circuit-manipulation studies suggest the cerebellum exerts direct influences on striatal reward processing (Chen et al., 2014) and dopaminergic tone in the ventral tegmental area (Carta et al., 2019). This architecture leaves the cerebellum well-positioned to integrate reward signals into ongoing cognitive control processes. Therefore, it is necessary to consider expanding existing *dual systems models* to better account for the role of the cerebellum in adolescent neurodevelopment. To address this need, we conducted a systematic search of the fMRI literature, synthesizing the current evidence on the cerebellum’s involvement in reward anticipation and feedback processing during adolescence.

### Cerebellar Circuit Anatomy

To integrate the cerebellum into models of adolescent neurodevelopment, we must first understand this region’s circuit makeup and functional involvement in reward processing (**Figure 1**). The physical architecture of the cerebellar circuit relies on relaying widespread neocortical data via a vast granule cell network, whose parallel fibers extend horizontally to synapse onto Purkinje cells where those signals intersect with sparse feedback from climbing fibers originating in the inferior olive of the medulla (Houck and Person, 2014; Tomasch, 1969). This structural convergence allows the Purkinje cells to rapidly adapt their synaptic weights, ultimately transforming high-dimensional information into the precise, timed release of deep cerebellar nuclei, which send excitatory outputs back to the neocortex via the thalamus (Diedrichsen and McDougle, 2026).

**Figure 1.**
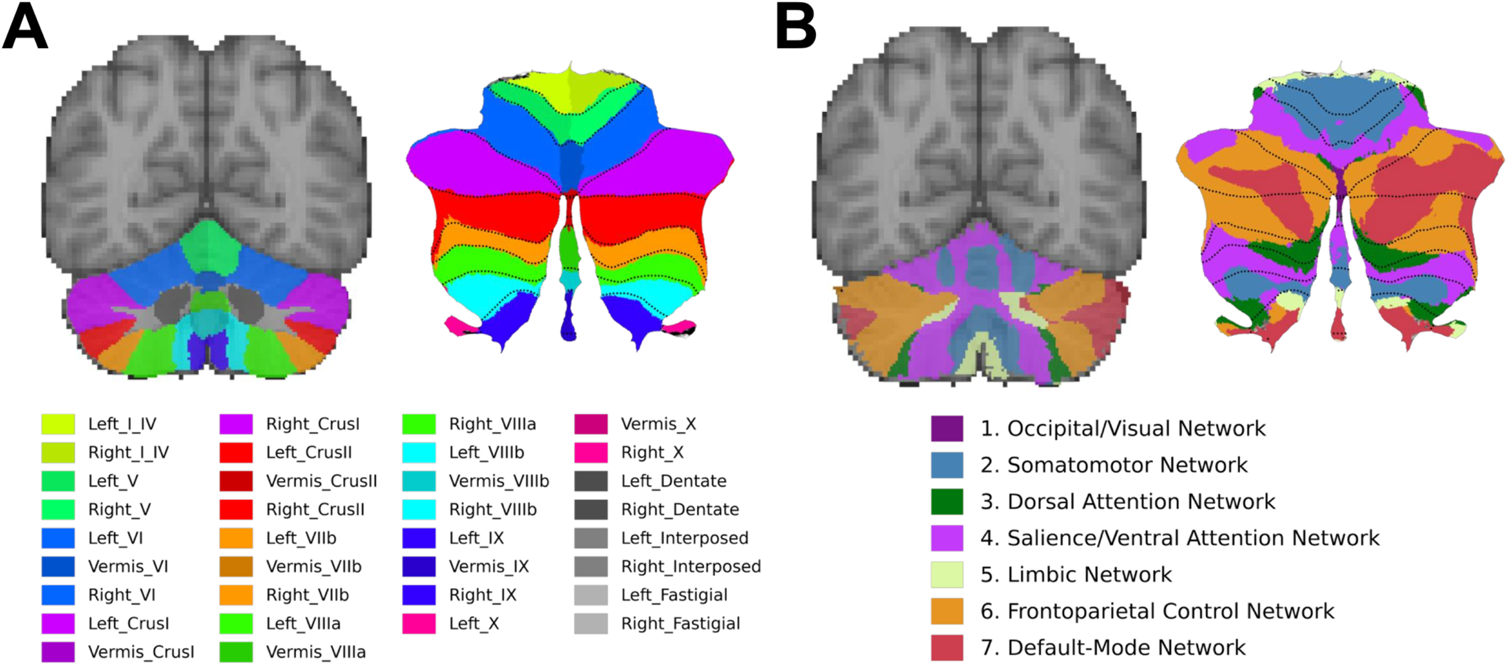
Anatomy and Functional Architecture of the Human Cerebellum. **(A)** Anatomical lobules of the human cerebellum, displayed on coronal slices alongside a 2D flatmap projection of the cerebellar cortex (Diedrichsen et al., 2009). **(B)** Cortico-cerebellar functional resting-state networks mapped onto the cerebellum. Note that the majority of the human cerebellum is densely and reciprocally connected to non-motor regions of parietal, prefrontal, and temporal cortices (Buckner et al., 2011).

The cerebellum is essential for motor control. The temporally-precise cerebellar circuit is thought to play a role in predicting and updating the sensory consequences of planned voluntary actions, ensuring that complex movements are smoothly executed and adjusted online to minimize errors (Wolpert et al., 1998). This ‘internal model’ function of the cerebellum is necessary in the sensorimotor domain, with extensive evidence that damage to this region causes impairments in the execution of complex movements (Kawato, 1999; Therrien and Bastian, 2015). However, converging lines of anatomical (Middleton and Strick, 1998), clinical (Schmahmann and Sherman, 1998), and functional imaging (Buckner et al., 2011; King et al., 2019) evidence have now firmly established that the cerebellum plays important functional roles in non-motor domains as well (for recent reviews, see (Diedrichsen and McDougle, 2026; Sokolov et al., 2017)). In fact, the majority of the human cerebellum is densely and reciprocally connected to regions of the parietal, prefrontal, and temporal cortices that contribute to a wide array of non-motoric cognitive and affective functions (Amore et al., 2021; Diedrichsen and McDougle, 2026; Schmahmann, 2019). The regions of the cerebellum that project to PFC, lobules Crus I and Crus II, appear to be particularly expanded structurally in humans relative to nonhuman primates, suggesting that these regions and their cortico-cerebellar circuit architecture may play a role in high-level cognition (Balsters et al., 2010). This new frontier in the science of the cerebellum includes strong evidence that it plays a functional role in reward processing (Kruithof et al., 2023).

### Cerebellar Reward Processing

Prominent among the non-motor domains that have recently been linked to cerebellar circuitry is the processing of rewards and the computations underlying reinforcement learning. In nonhuman animal model studies, the cerebellum has been shown to modulate dopaminergic tone in ventral tegmental area, and also forms a loop with the basal ganglia—projecting to striatum through the thalamus, and receiving return projections from striatum via the subthalamic and pontine nuclei (Bostan et al., 2010; Carta et al., 2019). At the circuit level, granule cells providing cortical inputs to the cerebellum appear to encode the expectation of rewards, whereas climbing fibers from inferior olivary nucleus to cerebellum encode post-feedback differences between what was expected and what was received (i.e., prediction errors; (Heffley and Hull, 2019; Kostadinov et al., 2019; Wagner et al., 2017)). These reward anticipation and feedback-locked computations are thought to contribute to error-driven learning and updating of an affective-motivational internal model, analogous to what the cerebellum contributes in the sensorimotor domain (Manto et al., 2024; Sokolov et al., 2017).

Rather than representing a redundant byproduct of cortico-striatal value computations, cerebellar reward signals appear to play a necessary functional role in reinforcement learning and decision-making. Indeed, patients with cerebellar degeneration show specific reinforcement learning deficits that highlight the cerebellum’s critical role in this credit assignment process (McDougle et al., 2016). Model-based fMRI in healthy volunteers suggests the cerebellum encodes an unsigned prediction error—indexing outcome ‘surprise’ or salience rather than a valenced reward value signal—following action execution (McDougle et al., 2019). These cerebellar reinforcement signals appear to be highly temporally sensitive. The region encodes not just the occurrence of an unexpected outcome, but the precise timing of that outcome relative to the internal forward model’s predictions (Trach et al., 2025). These cerebellar computations are reflected at the macro/functional brain network level in human neuroimaging studies, where several recent *meta*-analyses have identified cerebellar clusters in adult task fMRI studies involving reward anticipation and/or feedback via monetary incentives (Jauhar et al., 2021; Kruithof et al., 2023; Oldham et al., 2018; Wilson et al., 2018). To reconcile these macro-level findings with circuit-level computations, we propose that the expansive cortico-cerebellar network may help support reinforcement learning credit assignment by maintaining short-timescale *eligibility traces*—lingering neural representations of recent states or actions—via parallel fiber afferents. This allows subsequent climbing fiber prediction errors to precisely update internal forward models. Crucially, the timing of these cortico-cerebellar computations map onto established ‘proactive’ and ‘reactive’ goal-directed control strategies that are known to vary across development (Braver, 2012; Munakata et al., 2012)—the *proactive* control of decision-making via feedforward internal models during reward anticipation, versus the *reactive* maintenance of eligibility traces post-feedback. Here we integrate developmental research on reinforcement learning and proactive/reactive control as a central organizing lens for the results of the current ALE meta-analysis on cerebellar reward processing in adolescence.

Of special interest, a recent meta-analysis of fMRI studies on reward anticipation or feedback in adults focused exclusively on the cerebellum (Kruithof et al., 2023). This study found a diffuse set of clusters involved in reward anticipation, comprising several dorsal cerebellar lobules (lobules I-IV, V, VI), crus I, and anterior portions of the vermis (vermis VIIIa, VIIIb, and IX; (Kruithof et al., 2023)). For reward outcome, Kruithof et al. found a more focal set of clusters involved in feedback processing, primarily in the cerebellar vermis (vermis VI and vermis crus II) with some common activation in left lobule VI (Kruithof et al., 2023). From a mechanistic perspective, this functional divergence may reflect the distinct computational and physiological demands of these different reward processing stages. Specifically, while reward anticipation activity in cerebellum is likely driven by an established cortical prediction about the future outcome via dense parallel fiber inputs evoking a large BOLD response, reward outcome processing in cerebellum likely relies on more sparse reafferent climbing fiber teaching signals that produce a reduced BOLD response (Diedrichsen et al., 2024). Crucially, the majority of findings on cerebellum and its involvement in reward processing come from animal models or human adults. To fully integrate the cerebellum into existing models of adolescent neurodevelopment, a central question remains: What does the cerebellum contribute to reward processing in adolescence?

## Methods

### Overview: A Meta-Analysis of Adolescent Reward Processing and the Cerebellum

To fully integrate the cerebellum into models of adolescent neurodevelopment, we conducted a systematic literature search and coordinate-based fMRI activation likelihood estimation (ALE) meta-analysis of the adolescent functional neuroimaging literature on reward processing. Included studies were not restricted to those focused explicitly on the cerebellum, as many whole-brain analyses report relevant cerebellar coordinates despite few studies directly focusing on the region. We aimed to compare and contrast the results of the current adolescent meta-analytic analysis with Kruithof et al.’s recent meta-analysis of adult cerebellar fMRI studies on reward processing. The degree of overlap and/or dissociation between functional maps observed in the adult and adolescent literatures will allow us to adjudicate between three possibilities regarding adolescent neurodevelopment and the cerebellum:

I. MATURITY: If adolescent cerebellar activation during *both* reward anticipation and feedback closely mirrors adults, it suggests that cerebellar development does not likely impact the heightened reward-seeking characteristic of adolescence.
II. IMMATURITY: If adolescent cerebellar activation differs from adults during *both* reward anticipation and feedback, it suggests a generalized, ongoing maturation of cerebellar reward processing in adolescence.
III. PHASE-SPECIFIC MATURATION: Lastly, if *either* reward anticipation or feedback exhibits a selectively distinct pattern in adolescence, this would provide unique insight into the maturation of efferent versus re-afferent reward activity in the cerebellum.

Therefore, regardless of the findings in the current meta-analysis, understanding how they do or do not overlap with extant meta-analyses in adults will help to resolve whether and how the cerebellum should be integrated with existing neurodevelopmental frameworks of maturational imbalance in the adolescent brain.

### Study Selection

**Figure 2** depicts a PRISMA diagram of studies included in the current coordinate-based meta-analysis. To identify eligible studies, PubMed was searched with the following criteria: (“reward” OR “prediction error” OR “reinforcement learning”) AND (fmri OR “functional magnetic resonance imaging”) AND (development OR adolesc* OR child*) AND (healthy OR typical OR trajectory). This identified N=613 total records when the search was conducted in Spring 2024. Google Scholar was also searched, identifying 10 unique records that were not on PubMed. The search was then filtered to human articles, in English with population 10-18 years. Studies were excluded if they did not have event-related task fMRI designs, did not include task contrasts to isolate reward anticipation or feedback, did not include a group of neurologically-healthy subjects, or did not report at least one cluster with peak activation coordinates within the cerebellum.

**Figure 2.**
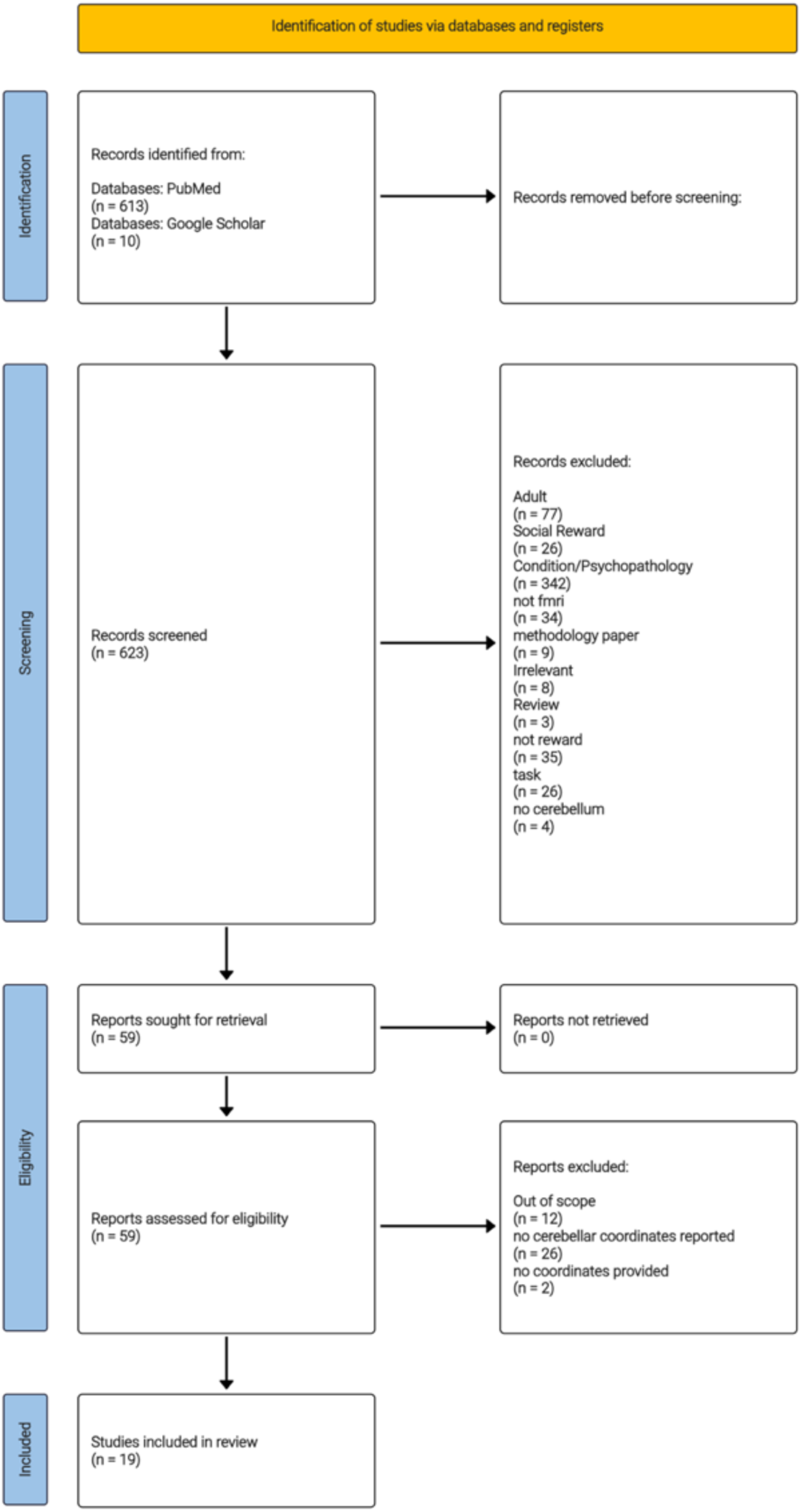
PRISMA diagram for study inclusion. The systematic literature search identified 623 initial records. After screening titles and abstracts against predefined exclusion criteria, 59 full-text reports were assessed for eligibility. 40 were excluded due to missing cerebellar coordinates or falling out of scope. A final total of 19 studies were included in the current meta-analysis.

### Activation Likelihood Estimation (ALE)

To determine the spatial convergence of cerebellar activations across studies, we conducted a coordinate-based meta-analysis using the Activation Likelihood Estimation (ALE) approach implemented in GingerALE (Eickhoff et al., 2009). Standardized coordinates reported in MNI space were retained as published. For studies originally reporting activation peaks in Talairach space, coordinates were converted to MNI space using the *tal2icbm* transformation integrated within GingerALE (Lancaster et al., 2007). Once all peaks were in a common MNI space, ALE fits the spatial uncertainty of each reported cluster peak across a collection of studies as a Gaussian probability distribution. This spatial uncertainty map is differentially weighted based on sample size to provide a robust meta-analytic statistical threshold for identifying convergent activation across studies. This ALE framework allowed us to directly compare and contrast cerebellar reward topologies in the current sample of adolescent fMRI studies to a recent adult ALE meta-analysis (Kruithof et al., 2023). Statistical significance for the resulting ALE meta-analytic maps was determined using a height threshold at *p*<0.001, and permutation-based cluster-extent thresholding with 1000 permutations to achieve a familywise error (FWE) rate at *p*_FWE_<0.05. The resulting meta-analytic maps were then projected onto a standard cerebellar anatomical atlas (Diedrichsen et al., 2009) using a flat map to directly visualize the degree of overlap or divergence from Kruithof et al.’s recent adult meta-analysis.

## Results

Initially, 613 total records were identified through PubMed with 10 additional unique records identified via Google Scholar (*N*=623 total studies). Title and abstract screening excluded 564 articles that were not within the scope of our systematic search. Fifty-nine reports were full text screened for inclusion and 19 articles met the current inclusion criteria (**Figure 2**). Overall, our ALE results reveal a spatial convergence across studies that is broadly consistent with a reward phase-specific developmental dissociation relative to mature functional topologies in adults (Kruithof et al., 2023)—with more focal cerebellar Vermis activation during reward anticipation, alongside more widespread activation of cognitive and affective cerebellar regions during feedback processing in adolescence (**Figure 3**).

**Figure 3.**
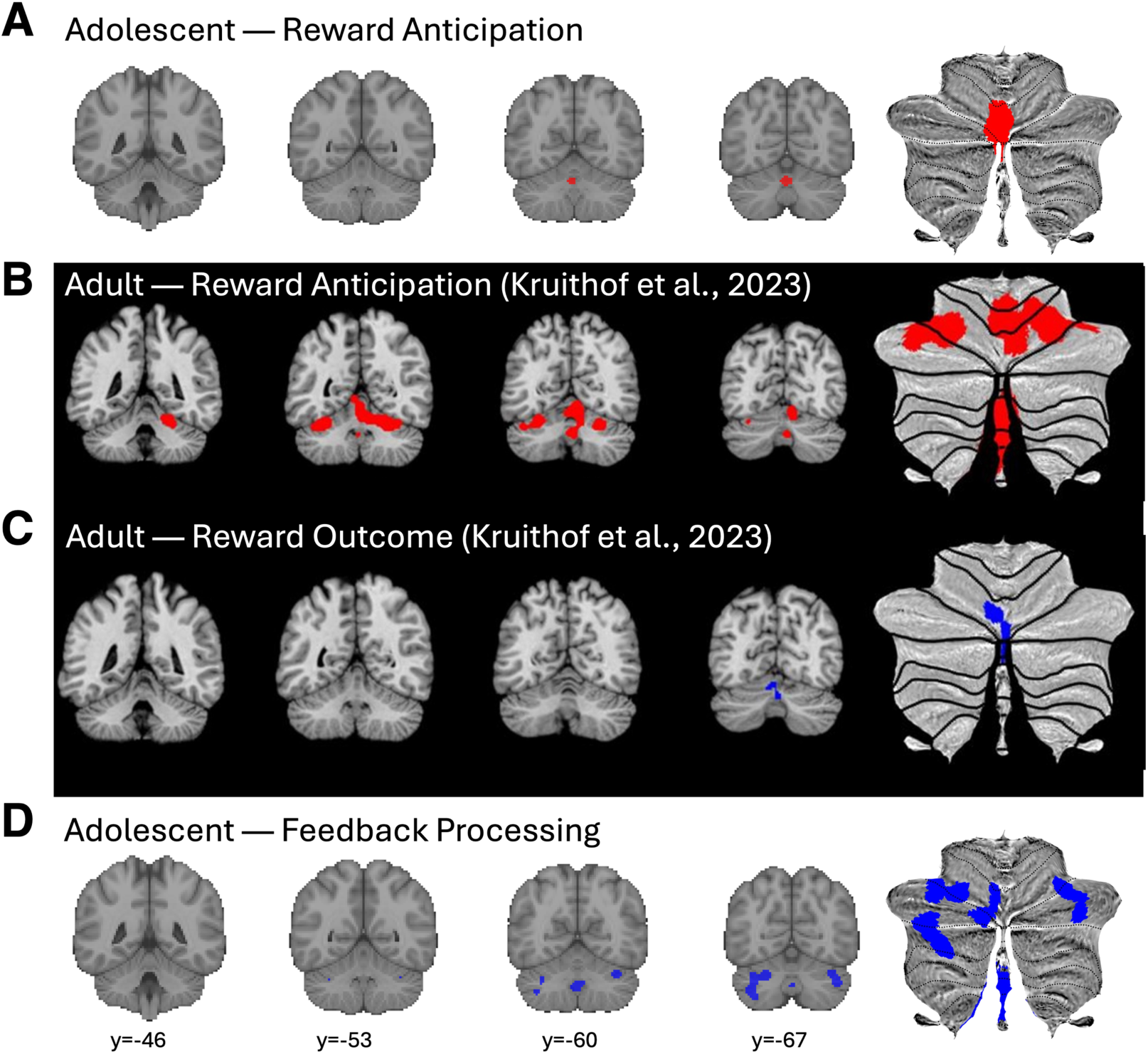
Developmental differences in cerebellar reward processing. **(A)** In the current ALE meta-analysis, adolescent reward anticipation elicited highly focal activation restricted primarily to the medial Vermis VI, contrasting with **(B)** the widespread, diffuse reward anticipation activity observed in adults (Kruithof et al., 2023). Conversely, **(C)** during reward feedback, adults exhibited a highly focal pattern primarily within the cerebellar vermis (Kruithof et al., 2023), while **(D)** the current meta-analysis found evidence for widespread, diffuse activation across cognitive and affective regions of the cerebellum (e.g., bilateral Crus I/II) during feedback processing in adolescents. Statistically thresholded ALE meta-analytic maps are depicted on coronal slices in MNI space, and projected onto SUIT flatmaps.

### Reward anticipation

A total of *k*=18 cerebellar clusters across *N*=7 studies were used in the reward anticipation ALE analysis (**Table 1**). One cerebellar cluster with two peaks was identified by the ALE of reward anticipation in adolescence (*z*≥4.26, *p*_FWE_≤0.022; **Table 2**). The peaks were located in Cerebellar Vermis VI and spread into adjacent regions of Lobule VI in the left hemisphere [(Diedrichsen et al., 2009); **Figure 1A**,**3A**]. These clusters overlapped most significantly with the regions of the cerebellum that are functionally connected to the cortical occipital and salience/ventral attention networks [(Buckner et al., 2011); **Figure 1B**,**3A**]. Notably, when compared to Kruithof et al. (2023), the cerebellar clusters observed in the reward anticipation contrast in adolescence most closely resembled those involved in feedback processing in adults (e.g. both were mostly restricted to the medial Vermis; **Figure 3**).

**Table 1.** List of included cerebellar clusters in the current coordinate-based fMRI meta-analysis. Full study list is included in Supplemental References.

| First Author | Contrast | Clust | Task | Age<sup>1</sup> | N | Peak MNI Coordinate |  |  |
| --- | --- | --- | --- | --- | --- | --- | --- | --- |
|  |  |  |  |  |  | X | Y | Z |
| Reward Anticipation |  |  |  |  |  |  |  |  |
| Bjork | Loss v. None | 1 | MID<sup>2</sup> | 12 to 17 | 24 | -75 | -46 | 4 |
| Forbes | Anticipation Early Adolescence | 1 | card guessing | $M\approx 11$ | 77 | -7 | -41 | -5 |
| Forbes | Anticipation Mid Adolescence | 1 | card guessing | $M\approx 11$ | 77 | -39 | -5 | -5 |
| Kappel | Gain v. Neutral | 1 | Incentive delay (child version) | $M\approx 11$ | 10 | 4 | -66 | -27 |
| Kappel | Gain v. Neutral | 2 | Incentive delay (child version) | $M\approx 11$ | 10 | -30 | -66 | 18 |
| Lorenz | Gain v. No Gain | 1 | Slot machine task | 13 to 16 | 34 | -2 | 0 | -55 |
| Nees | Big v. None | 1 | MID | $Mean\approx 14$ | 54 | -61 | -26 | -3 |
| Nees | Big v. Small | 1 | MID | $Mean\approx 14$ | 54 | -5 | -58 | 3 |
| Nees | Small v. None | 1 | MID | $Mean\approx 14$ | 54 | -3 | -70 | -26 |
| Nees | Anticipation Win | 1 | MID | $Mean\approx 14$ | 54 | -23 | -73 | 0 |
| Nees | Anticipation Loss | 1 | simple guessing | $Mean\approx 14$ | 54 | -16 | -82 | -6 |
| Nees | Anticipation Win | 1 | simple guessing | $Mean\approx 14$ | 54 | -76 | -22 | -2 |
| Telzer | Pumps v. Control | 1 | BART<sup>3</sup> | 14 to 16.5 | 48 | -50 | -26 | 26 |
| Telzer | Pumps v. Control | 2 | BART | 14 to 16.5 | 48 | -36 | -56 | -26 |
| Waltmann | Negative Choice Probability | 1 | Probabilistic reversal learning task | $M\approx 14.8$ | 40 | -32 | -38 | -56 |
| Waltmann | Negative Choice Probability | 2 | Probabilistic reversal learning task | $M\approx 14.8$ | 40 | -38 | -52 | -44 |
| Waltmann | Negative Choice Probability | 3 | Probabilistic reversal learning task | $M\approx 14.8$ | 40 | -30 | -56 | 36 |
| Waltmann | Negative Choice Probability | 4 | Probabilistic reversal learning task | $M\approx 14.8$ | 40 | -78 | -26 | -8 |
| Feedback Processing |  |  |  |  |  |  |  |  |
| Cao | Time 1 | 1 | MID variant | $M\approx 14$ | 13<br>0 | -49 | -20 | 0 |
| Cao | Time 1 | 2 | MID variant | $M\approx 14$ | 13<br>0 | -23 | -52 | 24 |

<sup>1</sup> Participant ages are presented as ranges whenever this information was included in the original manuscript. When the age range was not reported, we include mean age instead (*M*).<sup>2</sup> Monetary Incentive Delay (MID) task<sup>3</sup> Balloon Analogue Reasoning Task (BART)

|  |  |  |  |  |  |  |  |  |
| --- | --- | --- | --- | --- | --- | --- | --- | --- |
| Cao | Time 2 | 1 | MID variant | $M \approx 19$ | 12<br>4 | -21 | -55 | -20 |
| Cao | Time 2 | 2 | MID variant | $M \approx 19$ | 12<br>4 | -82 | -17 | 15 |
| Crowley | Decision Making | 1 | Colorado balloon game | 14 to 18 | 20 | -14 | -52 | -48 |
| Crowley | Decision Making | 2 | Colorado balloon game | 14 to 18 | 20 | -36 | -60 | 6 |
| Crowley | Decision Making | 3 | Colorado balloon game | 14 to 18 | 20 | -38 | -64 | 2 |
| Crowley | Decision Making | 4 | Colorado balloon game | 14 to 18 | 20 | -34 | -64 | -38 |
| Crowley | Wins | 1 | Colorado balloon game | 14 to 18 | 20 | -68 | -44 | -40 |
| Crowley | Wins | 2 | Colorado balloon game | 14 to 18 | 20 | -66 | -28 | 38 |
| Ernst | Win v. No Win | 1 | Wheel of fortune | 9 to 17 | 16 | -78 | -16 | 18 |
| Harle | High-Stake v. Baseline | 1 | gambling task with high and low stakes | 13 to 17 | 42 | -11 | -71 | -9 |
| Hawes | Gain table s3 | 1 | incentivized antisaccade task | 10 to 18 | 13<br>9 | -80 | 28 | -24 |
| Helfinstein | Contingent Pos. FB v. Neg FB | 1 | reward | 14 to 18 | 32 | -72 | -24 | 11 |
| Helfinstein | Contingent Pos. FB v. Neg FB | 2 | reward | 14 to 18 | 32 | -46 | -41 | -26 |
| Insel | Gain Greater than Loss | 1 | magnitude tracking task with low and high stakes condition | 13 to 18 | 74 | -66 | -38 | 42 |
| Insel | High Greater Than Loss Gain | 1 | magnitude tracking task with low and high stakes condition | 13 to 18 | 74 | -34 | -68 | -50 |
| Insel | High Greater Than Loss Gain | 2 | magnitude tracking task with low and high stakes condition | 13 to 18 | 74 | -24 | -66 | -30 |
| Insel | High Greater Than Low Gain | 1 | magnitude tracking task with low and high stakes condition | 13 to 18 | 74 | -24 | -66 | -30 |
| Insel | High Greater Than Low Gain | 2 | magnitude tracking task with low and high stakes condition | 13 to 18 | 74 | -28 | -62 | 36 |
| Nees | Win | 1 | MID | $M \approx 14$ | 54 | -61 | -41 | -3 |
| Nees | Loss | 1 | simple guessing | $M \approx 14$ | 54 | -58 | -40 | 0 |
| Nees | No Loss | 1 | simple guessing | $M \approx 14$ | 54 | -82 | -16 | -6 |
| Nees | No win | 1 | simple guessing | $M \approx 14$ | 54 | -14 | -78 | -6 |
| Nees | Win | 1 | simple guessing | $M \approx 14$ | 54 | -82 | -16 | -6 |
| Op de Macks | Boys: Reward Greater Than Loss | 1 | Jackpot Gambling Task | 10 to 16 | 50 | -12 | -72 | -3 |
| Vaidya | Amount Gain v. Failure Gain | 1 | modified MID | 12 to 15 | 18 | -36 | -66 | 42 |
| Vaidya | Amount Gain v. Failure Gain | 2 | modified MID | 12 to 15 | 18 | -36 | -68 | -34 |
| Van Leijenhorst | Reward 14-15yo | 1 | Slot machine task | 10 to 15 | 18 | -15 | -63 | 0 |
| Waltmann | Negative DU PES<sup>4</sup> | 1 | Probabilistic reversal learning task | $M \approx 14.8$ | 40 | -28 | -10 | -80 |
| Waltmann | Negative DU PES | 2 | Probabilistic reversal learning task | $M \approx 14.8$ | 40 | -36 | -64 | -28 |
| Waltmann | Negative DU PES | 3 | Probabilistic reversal learning task | $M \approx 14.8$ | 40 | -38 | -62 | -46 |
| Waltmann | Negative DU PES | 4 | Probabilistic reversal learning task | $M \approx 14.8$ | 40 | -32 | -72 | -50 |
| Waltmann | Positive Choice Probability < 50 | 1 | Probabilistic reversal learning task | $M \approx 14.8$ | 40 | -34 | -32 | -56 |
| Waltmann | Positive Choice Probability < 50 | 2 | Probabilistic reversal learning task | $M \approx 14.8$ | 40 | -40 | -52 | -48 |
| Waltmann | Positive Choice Probability < 50 | 3 | Probabilistic reversal learning task | $M \approx 14.8$ | 40 | -12 | -78 | -28 |
| Waltmann | Positive Choice Probability < 50 | 4 | Probabilistic reversal learning task | $M \approx 14.8$ | 40 | -32 | -56 | 34 |

<sup>4</sup> DU PES = Double-Update Prediction Errors

**Table 2.**
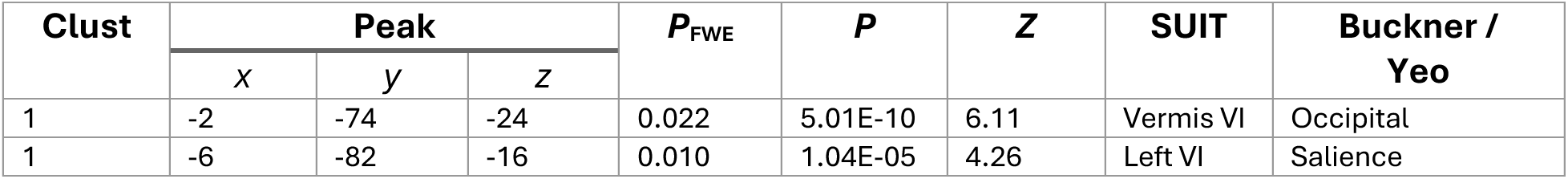
Full cluster and peak information from the adolescent reward anticipation ALE. Anatomical region-of-interest labels derived via cluster and peak overall with commonly used atlases (Buckner et al., 2011; Diedrichsen et al., 2009).

### Feedback processing

A total of *k*=37 cerebellar clusters across *N*=12 studies were used in the feedback ALE analysis (**Table 1**). Four cerebellar clusters with 10 peaks were identified by the ALE of feedback processing in adolescence (*z*≥3.76, *p*_FWE_≤0.025; **Table 3**). The peaks were located in Crus I bilaterally, Crus II in left hemisphere, left Lobule VI, and Vermis IX [(Diedrichsen et al., 2009); **Figure 1A**,**3D**]. These clusters overlapped with cerebellar regions that are functionally connected to the cortical frontoparietal, default mode, and limbic networks [(Buckner et al., 2011); **Figure 1B**,**3D**]. When compared to Kruithof et al. (2023), the diffuse cognitive and affective regions of the cerebellum recruited during feedback in adolescence more closely resembled those involved in anticipation reward in adults (e.g. both comprised Crus I/Crus II bilaterally; **Figure 3**).

**Table 3.** Full cluster and peak information from the adolescent feedback processing ALE. Anatomical region-of-interest labels derived via cluster and peak overall with commonly used atlases (Buckner et al., 2011; Diedrichsen et al., 2009).

| Clust | Peak | | | $P_{FWE}$ | $P$ | $Z$ | SUIT | Buckner / Yeo |
| --- | --- | --- | --- | --- | --- | --- | --- | --- |
| | $x$ | $y$ | $z$ | | | | | |
| 1 | -24 | -66 | -30 | 0.020 | 2.27E-08 | 5.47 | Left VI | Frontoparietal |
| 1 | -34 | -70 | -50 | 0.017 | 8.35E-07 | 4.79 | Left Crus II | Frontoparietal |
| 1 | -38 | -64 | -46 | 0.014 | 5.03E-06 | 4.41 | Left Crus II | Frontoparietal |
| 1 | -36 | -66 | -34 | 0.014 | 9.10E-06 | 4.28 | Left Crus I | Frontoparietal |
| 1 | -32 | -58 | -34 | 0.010 | 8.65E-05 | 3.76 | Left VI | Frontoparietal |
| 2 | -6 | -80 | -16 | 0.025 | 5.74E-10 | 6.09 | Left VI | Frontoparietal |
| 2 | -10 | -80 | -28 | 0.017 | 7.97E-07 | 4.80 | Left Crus I | Frontoparietal |
| 3 | 42 | -66 | -36 | 0.018 | 3.29E-07 | 4.97 | Right Crus I | Default-Mode |
| 3 | 36 | -62 | -28 | 0.016 | 1.47E-06 | 4.68 | Right Crus I | Frontoparietal |
| 4 | 0 | -60 | -40 | 0.019 | 8.66E-08 | 5.23 | Vermis IX | Limbic |

## Discussion

The prevailing neurodevelopmental models for understanding risky, reward-guided decision-making in adolescence center on a maturational imbalance between prefrontal cognitive control and motivational brain regions (Casey, 2015; Luna et al., 2015). While the current meta-analysis does not diminish these essential models, it highlights the need to incorporate cerebellar maturation into this framework. The cerebellum demonstrates a protracted development similar to the prefrontal cortex, alongside strong reciprocal connectivity with the specific prefrontal and striatal regions involved in reward-guided decision-making (Balsters et al., 2010; Bostan et al., 2010; Buckner et al., 2011; Tiemeier et al., 2010). By synthesizing 19 task fMRI studies in an ALE meta-analysis, the current results allow us to adjudicate between competing models of cerebellar development, and provide evidence for a reward phase-specific difference in cerebellar spatial topology during adolescence that is consistent with ongoing maturation. Specifically, the spatially diffuse BOLD responses in Crus I/II and Lobule VI regions of the cerebellum during feedback processing in adolescence—alongside circumscribed Vermal activations during reward anticipation—differs markedly from prior findings in adults (Kruithof et al., 2023), suggesting a potentially important functional dissociation in the adolescent brain. Here, we propose that this widespread cerebellar recruitment during feedback processing in adolescents—but not adults— could capture the distinct computational demands of reinforcement learning during this developmental window.

### Cerebellar BOLD and Reward-Based Internal Models

Before contextualizing the current results in the framework of adolescent reinforcement learning, the spatial extent of the BOLD signal should be contextualized through the structural constraints of cerebellar microcircuitry. The magnitude and spatial extent of the BOLD signal are fundamentally constrained in cerebellum. Climbing fibers, which originate in the inferior olive, act as discrete ‘teaching’ signals that encode reward prediction errors; while they powerfully depolarize Purkinje cells via complex spikes, they fire at extremely low rates (Raymond and Medina, 2018). In contrast, high-dimensional contextual information arriving to the cerebellum from the cortex via dense parallel fiber networks tend to fire at higher rates than the climbing fiber pathways (Wagner et al., 2017). These circuit-level differences might contribute to distinct metabolic demands in the cerebellum. Specifically, the fMRI BOLD signal is primarily thought to reflect efferent signals from the neocortex arriving as dense parallel fiber afferents, rather than sparse re-afferent teaching signals arriving from the brainstem via the climbing fiber path (Diedrichsen et al., 2024). Accordingly, the BOLD response during reward anticipation *and* receipt are both more likely to scale with the magnitude and spatial topology of cortico-cerebellar connections, rather than the climbing fiber re-afferents.

In adult studies, reward anticipation is reliably associated with bilateral recruitment of cerebellar Crus I and Lobules I-VI, while feedback processing is associated with more circumscribed activation restricted mostly to the Vermis (Kruithof et al., 2023). In line with the interpretation Kruithof et al. (2023) took regarding their ALE of adult cerebellar reward processing, we believe this pattern may be driven by the predictive nature of reward-guided decision-making in adults. During reinforcement learning (RL) in adults, internal forward models of task contingencies are learned rapidly and stabilize quickly. For example, adults are better able to track temporal dependencies to learn underlying relational task structures, across longer time horizons suggesting longer retention of eligibility traces, and are also able to engage in counterfactual (i.e., latent) outcome-dependent learning (Nussenbaum et al., 2026; Palminteri et al., 2016). By combining this capacity for internal model construction, alongside reduced decision noise, adults can use feedforward information to more accurately predict likely choice outcomes [cf., (Chierchia et al., 2023; Master et al., 2020; Nussenbaum and Hartley, 2021; Xia et al., 2021)].

In the sensorimotor domain, internal forward models in cerebellum support adaptive, feedforward motor control (Ito, 2008; Wolpert et al., 1998). It is plausible that a similar predictive coding architecture may support model-based, reward-guided decisions. Specifically, widespread activation of cognitive and affective regions of the cerebellum during reward anticipation in adults (Kruithof et al., 2023) may reflect the activation of an internal reward-based model, which can then be used to compute a reward prediction error teaching signal to optimize future choices (Trach et al., 2026). Internal model construction in adults is known to engage common underlying cognitive resources as executive function and cognitive control (Otto et al., 2012). Since adult activation during reward anticipation is prominent in bilateral Crus I—a cerebellar region that has dense connectivity to the frontoparietal control network and is reliably activated in tasks with working memory and executive control demands (Buckner et al., 2011; Nettekoven et al., 2024)— this suggests that cortico-cerebellar circuitry may support ‘proactive’ cognitive control during motivated decision-making and RL tasks in adults.

### Proactive Versus Reactive Cerebellar Processing in Youth and Adults

In dual mechanisms of control (DMC) theory, ‘proactive’ control refers to the preparatory maintenance of task-relevant information in frontoparietal circuits to bias downstream cognitive processes and enable goal-pursuit, and is dissociable from ‘reactive’ cognitive control that is linked to ventral attention / salience network activity and is only transiently deployed in response to a salient stimulus after it has been presented (Braver, 2012). Several lines of research suggest proactive control is cognitively effortful and develops slowly, and that the ability to exert proactive control appears to be online by late childhood (Chatham et al., 2009; Munakata et al., 2012). While the *ability* to use proactive control being established in adolescence, the *propensity* to use it may be shaped by the recruitment of dorsolateral prefrontal cortex and inferior frontal junction regions of the frontoparietal network, which demonstrate marked changes in recruitment during proactive control between adolescence and early adulthood (Andrews-Hanna et al., 2011).

Based on the current ALE analysis, adolescents demonstrate a functional dissociation in the temporal pattern of cerebellar activations during reward anticipation and feedback relative to adults. Specifically, adolescent cerebellar activation during reward anticipation is highly focal and spatially restricted to the medial Vermis—in clusters that are functionally connected to the occipital and salience networks. Reward has a robust behavioral and neural impact on bottom-up, exogenous attention (Ivanov et al., 2012; Le Pelley et al., 2022), and this might be what drives activation in these regions during reward anticipation in adolescents. In contrast, feedback processing in the current ALE analysis was associated with extensive bilateral recruitment of Crus I/II, Lobule VI, and Vermis IX. These feedback-locked activations included regions that are most strongly connected to the brain’s frontoparietal control network, and task-based studies suggest these regions span multiple task-based domains, including those involved in complex, cognitively demanding processes like working memory maintenance and divided attention (Buckner et al., 2011; King et al., 2019; Nettekoven et al., 2024).

The current results may represent an analogue to this proactive versus reactive developmental shift in the domain of reward. Specifically, the current ALE may indicate that adolescents may not typically form complex internal models during reward anticipation to generate feedforward predictions about upcoming rewards. Instead, reward anticipation in the adolescent cerebellum may be more associated with maintaining basic arousal and visual attention processes. In contrast, more computationally demanding processing of rewards in frontoparietal and limbic networks may be done *reactively* in adolescence, in the post-reward feedback period.

These findings fit with models of adolescent reward-based learning or RL, which suggest that youth disproportionately rely on recent information to guide motivated choice behavior. Specifically, adolescent expectations appear to be primarily shaped by recent outcomes, and only those observed via direct experience without the capacity for counterfactual RL (Nussenbaum et al., 2026; Nussenbaum and Hartley, 2019; Palminteri et al., 2016). The current ALE findings suggests reactive control of behavior may be the dominant cognitive strategy deployed by adolescents during reward processing tasks. Ultimately, this may drive more decision noise in reward-guided choices in adolescents relative to adults (Chierchia et al., 2023; Xia et al., 2021).

Notably, the current findings align closely with recent cross-species literature on the cerebellar substrates of reward processing. In both rodents and nonhuman primates (NHPs), lateral cerebellar regions appear to play a critical role in feedback processing and behavioral monitoring (Heffley and Hull, 2019; Kostadinov et al., 2019; Sendhilnathan et al., 2020; Wagner et al., 2017). Of special interest to the current ALE results, a recent NHP study found that Purkinje cells in Crus I and Crus II of the cerebellum carry prediction error-related signals that drive reinforcement learning and internal model updating during a reward-based two-alternative forced-choice task (Sendhilnathan et al., 2020). In the human neuroscience literature, using fMRI BOLD imaging in adult participants, Crus I/II and Lobule VI appear to track temporally-sensitive reinforcement learning signals, and interact functionally with the striatum during reward feedback (Lee et al., 2025; Trach et al., 2026). The current results converge with these findings. They suggest that during adolescence—a life stage where learning mechanisms are *acutely* sensitive to recent outcomes (Nussenbaum and Hartley, 2019)—Crus I/II and Lobule VI are highly responsive during feedback processing.

### Limitations

Importantly, our approach—comparing results from a prior adult cerebellum ALE meta-analysis (Kruithof et al., 2023) with our own *de novo* ALE on adolescent studies—should be interpreted with caution. Given the relatively limited number of adolescent reward processing fMRI studies meeting inclusion criteria, it was necessary to include a broader array of experimental designs to adequately capture reward anticipation and feedback processing. Consequently, the resulting list of studies exhibits a greater degree of task heterogeneity compared to the adult ALE, which relied on the monetary incentive delay (MID) task variant for 77% of its included studies (relative to 35% of studies in the current adolescent meta-analysis). Therefore, this increase in task heterogeneity may introduce spatial variance in the current developmental findings relative to the more homogeneous tasks integrated into the prior adult ALE (Kruithof et al., 2023).

Potential cross-sectional differences may also exist in field standard methods in adult versus pediatric fMRI studies of reward processing. Due to the difficulties inherent to pediatric task fMRI studies, for example, it is possible that there would be systematic differences in scan duration (e.g., reduced trial counts, variances in trial timing or event jitter), and/or in fMRI head motion-related confounds (e.g., higher mean framewise displacement) in youth relative to adult studies. Furthermore, cerebellar reinforcement signals are highly temporally sensitive, meaning that variances in feedback delays between adult and pediatric paradigms could theoretically influence BOLD signal magnitudes [cf., (Diedrichsen and McDougle, 2026)]. Therefore, it is possible that some apparent differences in meta-analytic cerebellar activations may be driven by these cohort differences in fMRI signal quality rather than representing developmental differences in reward processing *per se*. These provide an important consideration for interpreting the present ALE results and their comparison to the prior adult ALE, but notably, the reward phase-specific nature of the observed spatial dissociation suggests that they may not fully account for the apparently contrasting results.

## Conclusions

The protracted maturation of the cerebellum occurs in parallel with the maturation of prefrontal cognitive control circuitry, and both are associated with profound changes in reward-guided behavior in youth. The current results suggest that existing fronto-striatal models of impulsive and risky reward-guided behavior in adolescence should be updated to incorporate the cerebellum. Importantly, while differences in sample size and spatial smoothing between adult and adolescent meta-analyses could partially contribute to the focal versus diffuse spatial topologies observed during feedback processing, the widespread adolescent activations reported here are not randomly distributed. Rather, they systematically map onto functionally relevant cortico-cerebellar networks. Thus, the widespread cerebellar activity seen during feedback processing in adolescence in the current meta-analysis may represent a more reactive form of reward processing in youth relative to adults. This may therefore represent an important stage in the normative development of more efficient, adaptive motivated decision-making in adulthood.

## ACKNOWLEDGMENTS

This work was supported by the National Science Foundation award number #2237795 and the National Institutes of Health (via National Institute on Alcohol Abuse and Alcoholism, award number (R01AA030283)). We greatly appreciate the constructive feedback of Drs. Lynette Abrams-Silva, Benjamin Clark, and Anila D’Mello on a draft of this manuscript during TSM’s PhD defense. We also thank Dr. James Cavanagh for his helpful comments on the manuscript.

